# Programmable 3D cell alignment of bioprinted tissue via soft robotic dynamic stimulation

**DOI:** 10.1101/2024.11.03.621771

**Authors:** Luca Rosalia, Katelyn Mosle, Soham Sinha, Yi Yi Du, Fredrik Solberg, Allison Jia, Jonathan Weiss, Tony Tam, Jessica E. Herrmann, Mark Cutkosky, Mark Skylar-Scott

## Abstract

Recent breakthroughs in biofabrication have enabled the development of engineered tissues for various organ systems, supporting applications in drug testing and regenerative medicine. However, current approaches do not allow for dynamic mechanical maturation of engineered tissue in 3D. Although uniaxial mechanostimulation techniques have shown promise in generating anisotropic tissues, they fail to recapitulate the biomechanics of complex tissues. As a result, existing biofabricated tissues lack the ability to replicate complex 3D alignment patterns essential for functional biomimicry. Here, we present a soft robotics-driven approach for programmable 3D alignment in 3D bioprinted tissue. Our method introduces the co-printing of biological tissue with a silicone-based soft robot via a custom core-double shell nozzle. The application of 3D, exogenous, dynamic expansion and torsional forces to the tissue via the co-printed silicone robot was found to drive cell alignment. Confocal imaging revealed pronounced anisotropy of the stimulated tissue samples compared to the unstimulated controls. In addition, different cellular orientation patterns resulted from each mode of stimulation, demonstrating the versatility of the soft robotic approach in tailoring the pattern of tissue alignment based on programmed mechanostimulation.

## Introduction

Advances in biofabrication have led to the development of more biologically accurate engineered tissue across a variety of organ systems. These advances have translated into better preclinical platforms for drug testing and have raised hope for regenerative medicine applications or transplantation [1]. However, current approaches do not allow for dynamic mechanical maturation of engineered tissue in 3D. As a result, to date, biofabricated tissues cannot mimic the complex 3D alignment of native tissues, hindering their clinical translation. Although 1D mechanostimulation approaches have led to the development of anisotropic tissues, such as skeletal muscle [2, 3], tendons and ligaments [4, 5], and cartilage [6], these techniques are not sufficient to engineer biomimetic tissues with 3D biomechanical function, such as the heart. In fact, the heart offers a notable example of how replicating the complex 3D alignment at both the cellular and tissue levels is crucial to achieving its physiologic function. In the native cardiac ventricle, myocardial cells are aligned along complex 3D helical patterns to maximize cardiac output [7]. Uniaxial cell contraction along these complex patterns generates shortening in both the circumferential and longitudinal directions, as well as twisting of the left ventricle.

Methods for inducing alignment in engineered cardiac constructs can be broadly divided into passive and active approaches. Passive techniques rely on anisotropic materials on which cardiomyocytes are seeded, while active stimulation involves the use of dynamic forces to promote tissue maturation [8]. Passive conditioning is often achieved by modulating substrate stiffness and topography or applying static stresses to cardiac tissues. For example, substrates like polyacrylamide gel or polydimethylsiloxane with elastic moduli of 10–20 kPa have been shown to promote cell alignment and increase cell spread area [9, 10]. Additionally, modifying substrate topography can enhance the anisotropic organization of cardiac tissue [9]. In 3D constructs, fiber-infused anisotropic gel scaffolds drive cell alignment in a 3D-printed ventricle [11]. Static stress facilitates sarcomere rearrangement and increases internal tension, advancing tissue maturation. For instance, one study demonstrated that maintaining hiPSC-CM-derived microtissues under static tension for two weeks using nylon tabs led to improved cell alignment, cardiomyocyte hypertrophy, enhanced contractility, increased passive stiffness, and better force-frequency relationships [12]. Similarly, static stresses applied to 3D-printed cardiac tissue grafts from hiPSC-CMs improved cell alignment, contractile force, extracellular matrix organization, and upregulated cardiac-specific gene expression [13]. Recently, bioprinted anisotropic building blocks of hiPSC-CMs have been developed by leveraging the shear forces experienced by the printed cells upon extrusion and programming anisotropy by varying the printed path [14]. Based on this approach, applied shear stress, stretching or extension force, and post-print deformation can be manipulated to create aligned structures in 3D cardiac tissues [15].

Active mechanical stimulation can further influence cardiomyocyte maturation, alignment, and contractile function [16, 17]. Uniaxial tensile forces are applied using mechanical, dielectric or pneumatic actuators [18–20]. Variations in these systems were developed to apply different regimes of uniaxial mechanical stimulation, including simple or cyclic uniaxial stretch or compression, for up to 8 days of stimulation [21]. In hiPSC-CM monolayers, contractile stress was found to increase with higher strain magnitudes, but reached a plateau at 15% strain. These studies supported that cyclic stimulation results in increased cellular elongation and enhanced physiological phenotypes [20–23]. Other studies have shown that active stimulation can support cardiomyocyte alignment in 3D constructs as well as 2D [24–26]. However, all of these studies limit their mechanical stimulation to uniaxial loading, which is not representative of the complex 3D mechanics of the native heart.

Here, we present a method to drive programmable 3D alignment of bioprinted tissue using tunable soft robots. This method combines the advantages of active mechanical stimulation and tailorable by programmably tuning the dynamic triaxial tissue strain. To produce 3D patterns of biohybrid constructs comprising tissue and silicone, we co-print biological tissue with a biocompatible, room-temperature vulcanizing silicone. The result is a tissue with an adhesive and integrated silicone soft robot. Pressurization of the silicone provides expanding forces to the cells, while twisting of the silicone-tissue construct is applied by a servo motor. Each of these 3D mechanostimulation modalities was found to result in a specific pattern of cellular orientation, as observed via confocal imaging. Although soft robots had already demonstrated their ability to recapitulate the motion and function of organ tissues in a physiological manner [27–33], this work demonstrates their potential application in 3D biomanufacturing and bioprinting to address complex tissue engineering challenges.

## Results and Discussion

### Workflow enabling co-printing and 3D mechanostimulation of silicone-tissue constructs

We first demonstrated the ability to co-print silicone and tissue as well as their mechanical coupling. A multi-material nozzle [34, 35] enabled the fabrication of the silicone-tissue-silicone filament illustrated in **Fig. 1a**. The mechanical behavior of the co-printed filament under tension is shown in Supplementary Movie 1. For this, we used commercial silicone (Dragon Skin 10, Smooth-On Inc.) and gelbrin, a composite of fibrinogen and gelatin. Then, we developed a specialized core-double shell nozzle that enables printing of 3D tissue-silicone coaxial tubular constructs (**Fig. 1b**). The nozzle was designed to extrude the bioink as the outer shell, silicone as the inner shell, and the gelatin microparticle suspension bath material as the core to support the silicone tube formation during the printing process. A low-cost 3D printer developed in our laboratory [35] was modified to enable extrusion of three materials mounted on a single z-axis (**Fig. 1c**, fig. S1). The coaxial printing occurs in 3 serial segments: a 15mm segment of silicone and core material, a 20mm segment of silicone, bioink, and core material, and another 15mm segment of silicone and core material (**Fig. 1c**). This core-shell construct is created at a speed of 50 mm/min in the z-axis to minimize disturbance to the supporting bath, resulting in a print time for one tube in approximately one minute (Supplementary Movie 2). A schematic of the final construct is visualized in **Fig. 1d**.

**Fig. 1.**
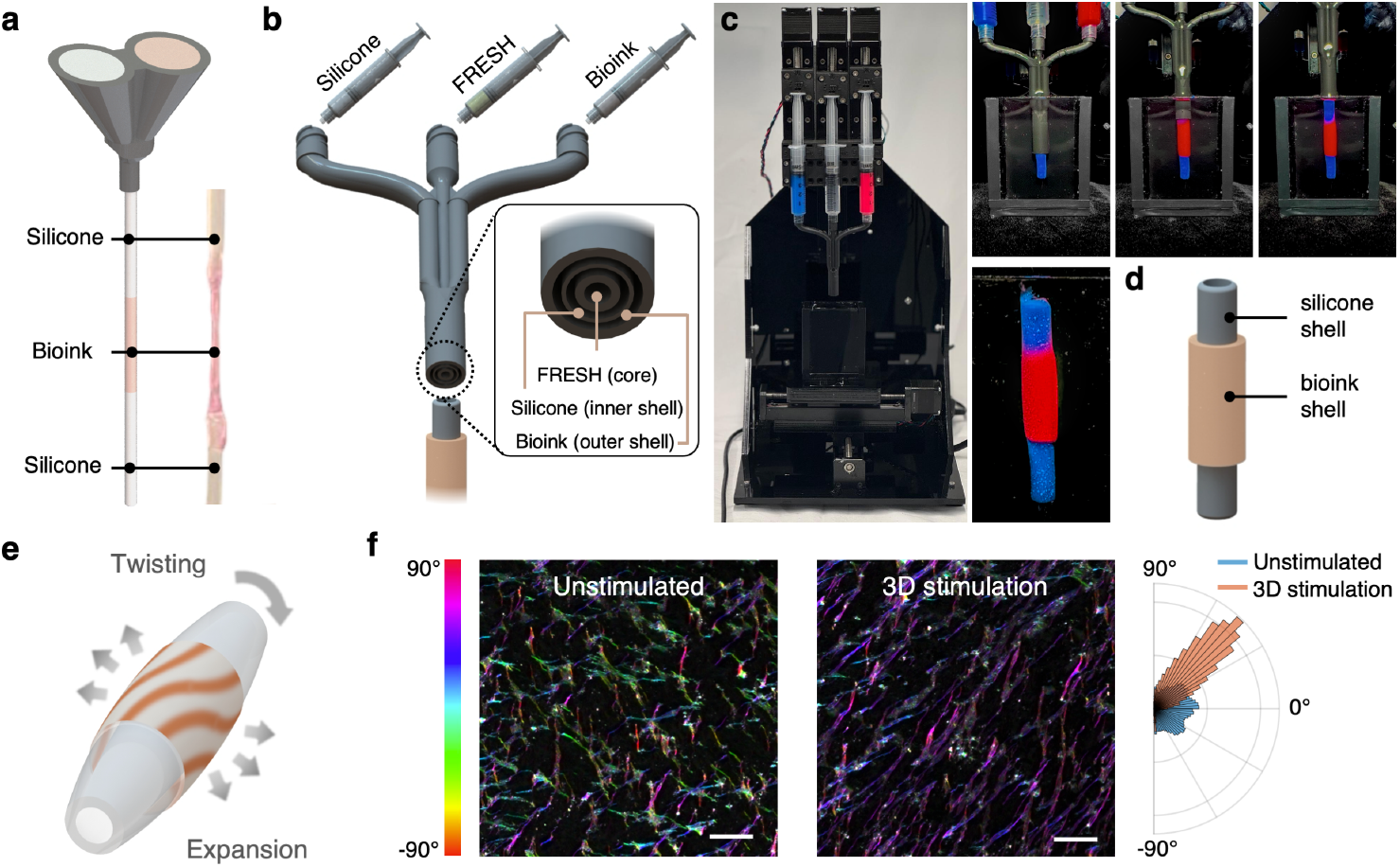
Silicone-tissue co-printing and 3D mechanostimulation workflow. **a**, Schematic and photography of silicone-bioink coprinting, highlighting mechanical adhesion between the two materials. **b**, Core-double shell nozzle for 3D printing of concentric and adherent silicone and tissue tubes. **c**, Photography of 3D printer and lapses of silicone-tissue co-printing. **d**, Schematic highlighting concentric silicone and tissue tubes after co-printing. **e**, Illustration of twisting and expansion regimes of 3D mechanostimulation enabled by silicone robot. **f**, Examples of unstimulated and stimulated tissue alignment on confocal imaging, highlighting anisotropy resulting from 3D mechanostimulation. Scale bars = 50 um.

We applied exogenous forces to tissue through the silicone robot to drive complex patterns of cellular and tissue alignment (**Fig. 1e**). Specifically, we applied torsion and expansion forces to the tissue in an active, dynamic, and cyclical manner, and visualized cellular elongation and alignment via actin staining and confocal microscopy. We characterized alignment resulting from various regimes of 3D mechanostimulation, including expansion only, twisting only, and both expansion and twisting. Results indicated that tissues subjected to active dynamic stimulation exhibited pronounced cell alignment and anisotropy compared to unstimulated tissue (**Fig. 1f**) and that the direction of alignment can be programmed by varying the mechanostimulation modality.

### Mechanical reactor design and characterization

In this work, we used a bioink made of neonatal human dermal fibroblasts (NHDF), herein referred to as fibroblasts (FBs), in a gelbrin ink. **Fig. 2a** outlines the bioprinting pipeline. Bioink preparation involved preheating the gelatin to 37°C, prior to adding fibrinogen to the centrifuged cell pellet. The gelatin was added to the fibrinogen-FB mixture and transferred to a syringe. To induce gelling and achieve rheological properties compatible with the surrounding FRESH support bath while also preventing cell settling, the syringe was cooled on ice. Frequent rotation of the syringe during this step ensured a homogenous cell distribution. Extrusion of the bioink into the FRESH (freeform reversible embedding of suspended hydrogels bioprinting) support bath containing thrombin (10 u / ml) prompted the polymerization of fibrinogen into fibrin. The printed construct was finally incubated at 37°C to allow the silicone to cure and melt the gelatin, resulting in a fibrin-cell construct co-printed around the silicone tube. High-throughput co-printing was achieved by replacing the syringes with new silicone and bioink materials. Switching to an acellular bioink in the syringe allowed us to print the residual bioink in the nozzle’s dead volume (**Fig. 2b**).

**Fig. 2.**
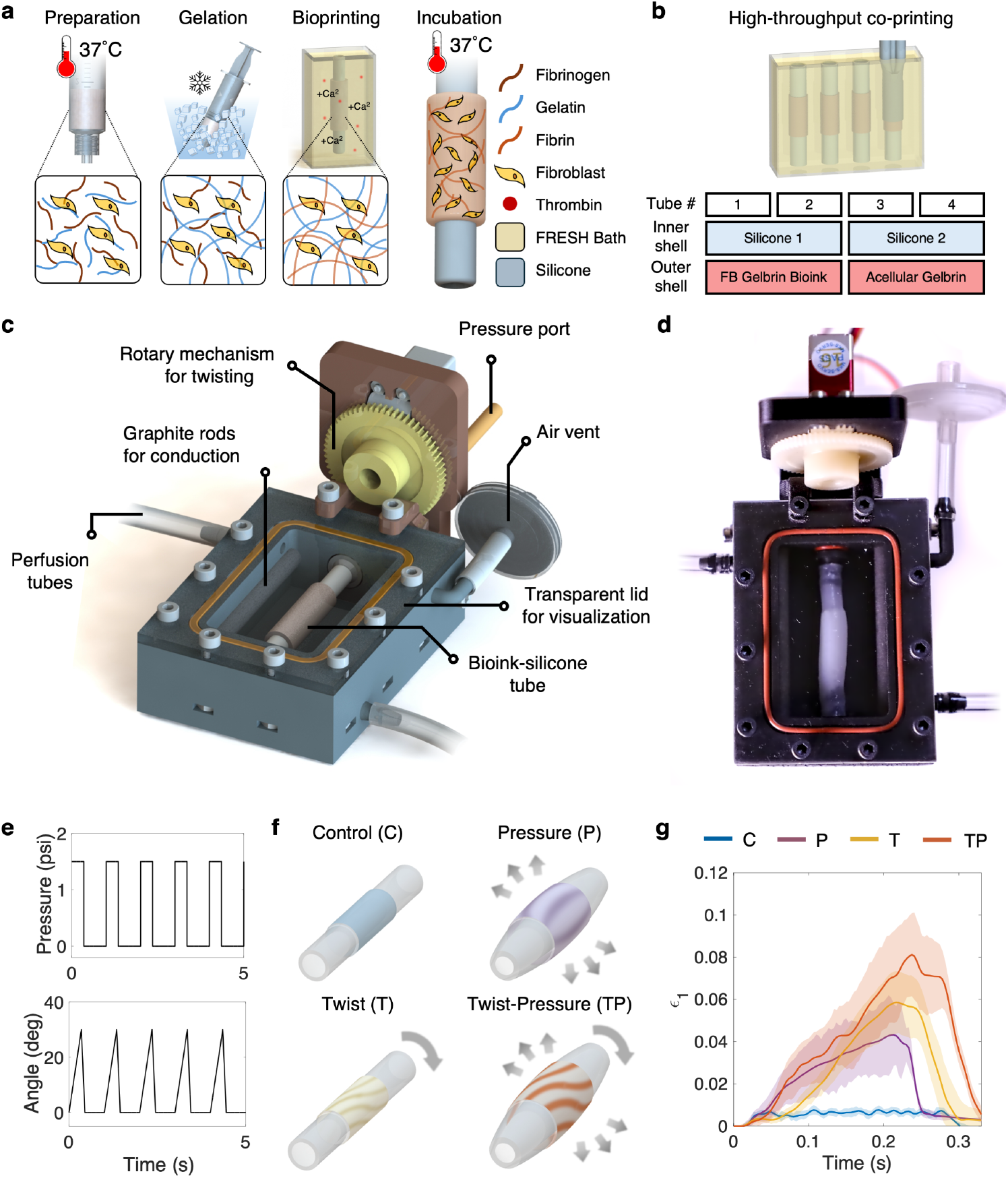
3D bioprinting and 3D mechanostimulation. **a**, Illustration of the 3D bio-printing steps in FRESH using our core-double shell nozzle. **b**, Schematic of high-throughput printing of silicone-tissue constructs. **c**, Schematic of mechanical reactor for 3D mechanostimulation. Pressure port and rotary mechanism enable simultaneous expansion of twisting of the tissue construct via the silicone robot. **d**, Image of the reactor with the bioink-silicone tube mounted. **e**, Input signals for the expansion and twisting of the constructs over five consecutive cycles. **f**, Illustration of 3D mechanostimulation modalities (control, twist, pressure, twist-pressure). **g**, Corresponding first principal strain measured via DIC. DIC graphs obtained at 3 psi expansion and 30° rotation.

The mechanical reactor was designed to provide 3D mechanical stimulation to the co-printed tissue-silicone constructs (**Fig. 2c-d**). A pressure port enables expansion of the constructs, while a rotary mechanism, consisting of a servo motor and two interconnected gears, allows for twisting of the samples via rotation of the rod onto which the samples are mounted. The pressure port is connected to a pressure regulator and solenoid valves that control the pressurization of the constructs. Graphite rods parallel to the tissue can enable electrical stimulation of the sample, if required. The reactor includes three ports; two for continuous perfusion and media exchange, and one for maintaining atmospheric pressure in the chamber, regardless of the pressurization state of the silicone tube. Detailed assembly instructions of the reactor are provided in fig. S2. Pressurization (1-2 psi) and twisting (30°) of the tissue are controlled by a custom-made printed circuit board (fig. S3). The samples were actuated cyclically at a frequency of 1 Hz and with a duty cycle of 33% (**Fig. 2e**).

Schematics of all actuation modalities are shown in **Fig. 2f**. We characterized the strains induced by each different regime and varying actuation levels using 3D digital image correlation (DIC), reporting changes in the first principal strains during actuation. **Fig. 2g** illustrates representative graphs for peak expansion pressures of 3 psi and twisting angles of 30°. At these levels, first principal strains of 0.0433 ± 0.0145, 0.0585 ± 0.0148, and 0.0812 ± 0.0197 were achieved for the pressure-only, twist-only, and pressure-twist conditions, respectively. First principal strain measurements for 0, 3, and 4 psi and 0, 20, 30, and 45°, and all combined conditions, are shown in fig. S4. Maximum strains of 0.01436 ± 0.014 were recorded during simultaneous expansion and twisting at 3 psi and 45°.

### 3D mechanostimulation induces programmable tissue anisotropy

Evaluation of tissue alignment showed pronounced tissue anisotropy due to 3D mechanostimulation compared to the unstimulated control samples. Representative images from n = 2 samples, each under four stimulation conditions (control, pressure, twist, twist-pressure) are shown in **Fig. 3a**. Qualitative analysis of F-actin and DAPI staining images, highlighting the actin cytoskeletal protein and cell nuclei, shows that both the control samples exhibit random patterns of cell orientation. Conversely, all three stimulated conditions show anisotropy, whereby the preferred cell orientation varies depending on the stimulation regime.

**Fig. 3.**
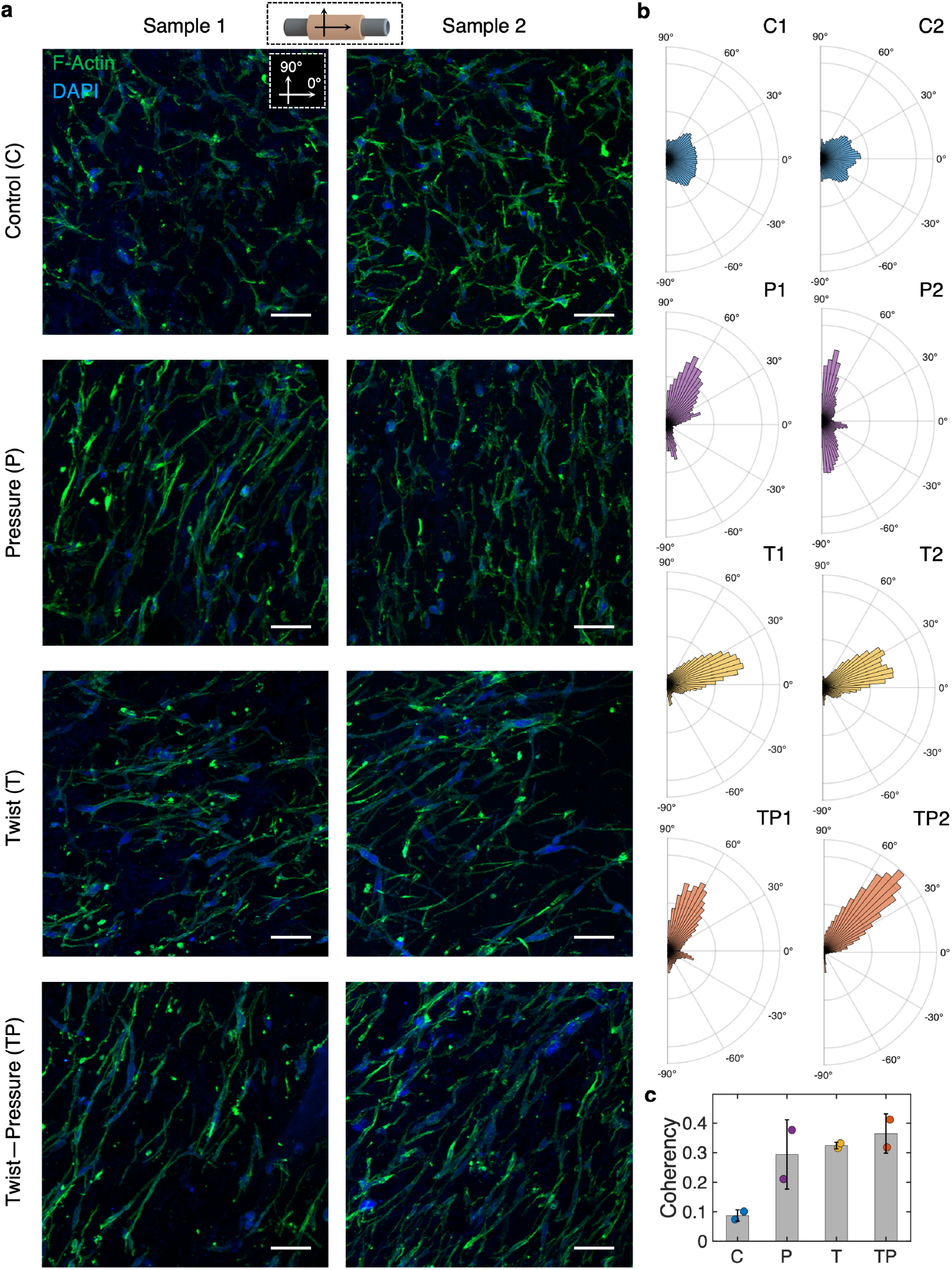
Tissue alignment resulting from 3D mechanostimulation. **a**, F-actin and DAPI stains of confocal images of tissues subjected to no mechanical stimulation (control, C), pressure only (P), twist only (T), and both twist and pressure (TP) from n = 2 biological replicates. **b**, Polar plots of cell alignment angle across all samples. **c**, Coherency plot of cell anisotropy. Scale bars = 50 um.

Quantitatively, polar histograms of the actin orientation obtained from these images highlight differences in the alignment angle (**Fig. 3b**). In all images, the 0° angle corresponds to the longitudinal axis of the tubular constructs. The average orientation angle between the two samples under pressure only was 75.2 ± 19.9°. This suggests that, in tubular structures under hoop stress, cells align circumferentially, which is consistent with alignment of smooth muscle cells in blood vessels [36]. Under twist-only mechanostimulation, the average alignment angle was 21.6 ± 5.1°. The samples subjected to both pressure and twisting exhibited an alignment angle of 52.7 ± 13.5°. Combination of these two actuation modalities results in helical stress patterns, which could be relevant to mimic physiologic alignment of cardiomyocytes for cardiac tissue engineering applications. In fact, in the native heart, cardiomyocytes are oriented along 3D helical patterns at an angle of about 50-60° [7, 37]. Our preliminary data suggest that these patterns of cellular orientation could be recapitulated using our proposed system. Evaluation of the coherency anisotropy further validates these findings (**Fig. 3c**). The lowest coherency was found to be associated with the unstimulated controls (0.0875 ± 0.0191), whereas combination of twisting and pressure resulted in the highest value (0.3650 ± 0.0665).

## Conclusions

This study presents the development of a soft robotics-driven approach for programmable 3D alignment in bioprinted tissues. By integrating a silicone-based soft robot with biological tissue using a custom core-double shell nozzle, we demonstrated that the application of dynamic, exogenous 3D forces can effectively promote cell alignment within the engineered tissues. Confocal imaging confirmed that the stimulated samples exhibited notable anisotropy compared to their unstimulated counterparts. In addition, the cellular orientation patterns seen with various mechanostimulation modalities indicate the versatility and promise of this approach in improving the functional mimicry of complex tissues. Particularly, pressurization of the constructs resulted in circumferential cellular alignment, whereas the combination of expansion and twisting led to angles analogous to those observed in native heart tissue. This research paves the way towards the development of biomimetic tissues with complex patterns of cellular alignment, bridging the gap between tissue engineering and pressing clinical challenges.

## Methods

### Mechanical reactor design

The mechanical reactor was designed to enable simultaneous mechanostimulation modalities of the 3D printed constructs. The base of the reactor and the rotating rod where the constructs are mounted were 3D printed on a Form 3+ stereolithography printer with biocompatible Biomed Black Resin (FP-F3P-01, Formlabs). The motor holder was 3D printed on a MarkTwo Carbon Fiber 3D printer (MF-M2-00, Markforged). All the 3D-printed components were designed in SolidWorks (2022, Dassault Syst`emes). The reactor lid was cut out of 1/8 inch-thick clear cast acrylic (8560K239, McMaster-Carr). A silicone o-ring (9319K158, McMaster-Carr) was added to create a watertight seal between the base of the reactor and the acrylic sheet.

The reactor features three ports; two for continuous-perfusion media exchange and one to maintain the pressure of the chamber to atmospheric level. All ports are connected to soft polyvinyl chloride tubing (5233K93, McMaster-Carr) via 3D-printed barbed connectors integrated with the base of the reactor.

Two 1/4-inches conductive graphite rods (9121K85, McMaster-Carr) are inserted bilaterally into the respective slots in the reactor and at an equal distance of about 2.5 cm from the edge of the tissue. The conductive rods allow for electrical stimulation of the tissue if needed. Alligator clips and conductive M2.5 screws are used to transmit current to the graphite rods.

Twisting is achieved by torque transmission from a servo motor (DS6125E, MSK servos USA) to the rotating rod via two gears; a 33-mm diameter gear (28106401, Maedler North America) attached to the motor and an 11-mm diameter gear attached to the rod (28102001, Maedler North America). A plastic bearing (6455K8, McMaster-Carr) and two o-rings (1173N01, McMaster-Carr) are attached to the rod to facilitate rotation while preventing leakage of the media out of the reactor.

Pneumatic pressure is delivered to the silicone-tissue constructs via a pressure port and through the rotating rod. The pressure level is adjusted through a digital pressure regulator (PCD-5PSIG-D-PCV10.30/5P, Alicat) and the airflow is controlled by a solenoid pressure manifold (VV5Q21-08N9FU0, SMC). Pressurization via the solenoid manifold and the twisting are synchronized via a custom printed circuit board (PCB).

### Board design and implementation

The PCB was designed to power and synchronize the modalities of mechanical stimulation of up to 12 mechanical reactors. The board allows for simultaneous actuation and independent control of each reactor, enabling dynamic stimulation under various conditions. The circuit consists of five ESP32 development boards (YEJMKJ): one for servo motor actuation, one for the solenoid valve, one for the cooling fans, and a master board. An additional board is used to enable biphasic electrical stimulation if needed.The board is connected to a single 32V-power supply (LW-K3010D, Longwei), drawing about 1 A at any given time. Each board was programmed on Arduino IDE (version 2.3.2).

### In vitro evaluation of construct mechanics

DIC was carried out on analogous cellular constructs under the same stimulation dynamics as used for mechanical training. The deformation of the tubes was measured for n = 1-3 tubes for a total of 11 distinct configurations, combining maximum angular rotation magnitudes of 0°, 30°, 45°, and 60° with inflation pressures of 0, 3, and 4 psi. 3D deformation was evaluated by measuring the first principal strain via DuoDIC, an open-source stereo DIC MATLAB toolbox [38]. 480 fps videos of samples mounted on a modified reactor were acquired simultaneously using two digital cameras (ZV-1, Sony) approximately at a 60° angle. The 3D space was then calibrated on the MATLAB’s Stereo Camera Calibrator App, using the frames acquired by the two cameras, and the 3D strains of the tissue were then reconstructed. The quality of the reconstruction was evaluated through the normalized least squares correlation criterion [39].

### Bioink preparation

The bioink was made of normal human dermal fibroblasts (NHDF), fibrinogen and gelatin. FBs were cultured in 2D flasks with high-glucose DMEM (11965126, Thermo Fisher Scientific) with 10% Fetal Bovine Serum (F4135, Sigma-Aldrich).

To prepare the bioink, the FBs were detached via a 3 minute 37°C incubation in TrypLE express enzyme (12605010, Thermo Fisher Scientific). The enzyme was then inhibited by adding FB medium (see above) at a 1:1 ratio with the enzyme. The FB flask contents were then transferred to 15 mL conical centrifuge tubes for centrifugation and cell counting via Trypan Blue assay (T10282, Invitrogen). The cell content was mixed in a 5 mL plastic syringe with final concentrations of 50 mg/mL of bovine fibrinogen (F8630, Sigma-Aldrich) and 7.5% type B gelatin (G6650, Sigma-Aldrich) in PBS++ with Calcium Chloride and Magnesium (14040117, Thermo Fisher Scientific) used for the dilutions. A volume of 2.5 mL was used for printing of four tubes, one per experimental group. The syringe with the bioink was cooled on ice for 5-10 minutes to allow it to gel before printing.

### Bath preparation

FRESH was prepared according to the published protocol [40]. A 50% v/v solution of ethanol and distilled water was heated to 45°C and stirred at 300 rpm. 2.0 wt% of gelatin type B (G7, Fisher Scientific), 0.2 wt% of pluronic F-127 (P2443, Millipore Sigma), and 0.1 wt% of gum arabic (G9752, Sigma-Aldrich) were added to the mix and dissolved at 45°C for 1 hr. The pH was then balanced to optimize for sphere dimensions (60-80 *μ*m). The pH level ranged between 5.2 and 7 based on the lot number of gelatin used. To achieve appropriate spherical formation of the gelatin, the mix was poured into 500 mL bottles and stirred at 250 rpm with an 8 cm stir bar overnight. The mix was then centrifuged at 500 g for three minutes and three additional PBS washes at 2000 g for two minutes before use or storage. FRESH was stored at 4°C in 50 mL conicals. 1% antibiotic-antimyotic (15240062, Thermo Fisher Scientific) was added for storage.

Before use, FRESH was centrifuged at 1500 g for 2 minutes. After the supernatant was removed, FRESH was poured into 80 mL flacktek cups and 10 units/mL bovine thrombin were added before mixing at 500 rpm for one minute. Subsequently, FRESH was transferred into a clear acrylic container for printing. A 5 mL syringe was loaded with FRESH to be extruded as the core material in our core-shell-shell nozzle for printing.

### Bioprinting and mounting on mechanical reactor

Three 5 mL syringes were loaded for printing and attached to the nozzle, containing FRESH as the core, silicone as the inner shell and bioink as the outer shell. Dragonskin 10 fast (Smooth-on) was used as the silicone, with the addition of 2 w% (of part A + B) of thinner and thickener agents (Smooth-on). The silicone syringe was centrifuged at 2000 g for 90 seconds for degassing.

A six-axis low-cost printer developed by our group was readapted to function with 3 extruders and one single z-axis [35]. Pronterface (Printrun version 2.0.1) was used as the user interface to control the printer. The code involved co-extruding the silicone (19.5 *μ*L/mm of z-axis movement), the tissue (25.8 *μ*L/mm), and the bath core (11.4 *μ*L/mm), while moving up with the z-axis to form a patent silicone-tissue construct. All motors moved at a rate of 50 mm/min. The constructs were printed to have a final length of 50 mm, consisting of a 20-mm tissue-silicone region flanked by two 15-mm silicone-only regions.

The printed tubes were left in the bath for 1 hour to ensure complete conversion of fibrinogen to fibrin by thrombin, before being transferred to individual 15 mL conical containers with medium and incubated overnight (Heracell Vios 250i, Thermo Fisher Scientific).

Each tube was then transferred to the reactor. The ends of the silicone tube were attached to the barbed connectors of the base of the reactor and the rod. Tissue adhesive (Vetbond, 3M) was added to secure the attachments. The tissue was kept hydrated by adding media drops via a transfer pipette through the mounting process every 30-60 seconds. After the tissue was mounted, the reactor was manually filled with medium. A hydrophilic filter was added at the inlet, and a hydrophobic filter was added at the air vent. The outlet port was connected via tubing to a drain bottle. The o-ring was then added into the reactor base and the acrylic lid was screwed on. The motor was then mounted on and secured with screws to the mechanical reactor and the pressure line of the solenoid valve was connected to the free end of the rod.

### Stimulation

Tissues were then dynamically stimulated for a total of seven days. The same parameters were used throughout the study: 30° torsion and 1-2 psi. A total of 8 constructs were divided into four groups and two biological replicates were carried out. The groups were defined by the modality of mechanical stimulation applied, namely pressure only, twisting only, both twisting and pressure, and none (control). All modalities of stimulation followed the physiologic time-varying elastance model [41], whereby actuation peaks at approximately 33% of the cardiac cycle. In this work, a constant stimulation frequency of 1 Hz or 60 bpm was used.

Full media changes were performed every 2-3 days using a 12-channel peristaltic pump (MFLX78006-24, Avantor) and 0.38 mm-pump tubing (MFLX95723-14, Avantor). Each set of pump tubes was connected to the mechanical reactor and the media reservoir via a combination of soft polyvinyl chloride tubing (5233K92, McMaster-Carr), 22G blunt needles (Nordson) and barbed connectors (51525K123, McMaster-Carr). The media reservoir was kept in a refrigerator at 4°C for the duration of the experiment. 1% antibiotic-antimyotic (15240062, Thermo Fisher Scientific) was added to the FB media as described above.

### Tissue staining

After cessation of the stimulation, the tissue was gently separated from the silicone tube using tweezers, and incubated in 4% paraformaldehyde (PFA) for 20 minutes after two PBS++ washes. Following fixation, samples were washed three times in PBS++ for 5 minutes each time. Prior to snap-freezing, tissues were placed in a 30% wt/v sucrose solution in PBS++ for 48 hours. Samples were then transferred to a 1:2 mix of 30% sucrose solution and optimal cutting temperature (OCT, 23-730-571, Fisher Scientific) for about 90 minutes. Each sample was then placed into a cryostat tissue mold, filled with 100% OCT, and frozen at -20°C using dry ice. Care was taken to ensure that all sucrose was removed. The samples were stored at -80°C prior to cryosectioning. The frozen tissues were sliced (30 *μ*m thickness) along the long axis using a cryostat (CM1950, Leica).

The tissue slides were thawed for 10-20 minutes at -20°C, followed by hydration in PBS++ for 10 minutes at room temperature. After aspiration of PBS, the tissue was incubated in a 1:1000 solution of Triton X (ab286840, Abcam) in animal free buffer AFB (1:5 with PBS ++) for 30 minutes at room temperature. The slides were washed twice with the AFB stock for 15 minutes each and a hydrophobic barrier was drawn around the sample using a PAP pen. The tissue was then incubated overnight at 4°C in an AFB solution with 1:200 Phalloidin (A12379, Thermo Fisher Scientific) and 1:1000 Hoechst 33342 (A3472, ApexBio). The slides were placed in a black-out box to prevent photobleaching. The tissue was then washed twice with the Tween-20 solution for 15 minutes each, followed by a single 15-minute wash with PBS++. Easy index (EI-100-1.52, LifeCanvas Technologies) was then added as a clearing solution. The slide was then covered with 22 x 30 mm #1.5 glass cover slip (CLS-1764-2230, Chemglass life sciences) before imaging.

### Structural evaluation

Samples were imaged on a confocal microscope (ZEISS LSM 980) and acquired using the Zen software (Zen 3.8, ZEISS). Lasers with wavelength 405 nm and 488 nm were used. Z-stack images were taken in full-Z stack per track mode, with a frame size of 512 x 512 px, and a frame time of about 2.0 sec and 8x averaging. Images were acquired at 2.5x, 10x, and 20x magnification. Alignment was processed using the OrientationJ ImageJ plug-in (version 1.54g) from 20x images. Alignment plots were graphed in Matlab (R2024a, MathWorks).

## Acknowledgments

Research reported in this publication was funded by the ARPA-H Health Enabling Advancements through Regenerative Tissue (HEART) Program (1AY1AX000002-01), Additional Ventures Cures Collaborative, and a Chan-Zuckerberg Biohub Investigator Award. L.R. was funded by the Burrough Wellcome Funds Career Awards at the Scientific Interface. S.S. and J.W. were funded by the National Science Foundation Graduate Research Fellowship Program. S.S was funded by the Stanford Graduate Fellowship. J.W. was funded by Stanford Bio-X. J.E.H. was supported by the Dorothy Dee and Marjorie Helene Boring Trust Research Award (Stanford Cardiovascular Institute) and the Berg Scholars Program (Stanford University School of Medicine).

## Author contributions

M.A.S.S, and L.R. designed the study. L.R., K.M. conducted the study, designed and built the mechanical reactor and the printing nozzle. L.R. performed staining, confocal imaging, and alignment analysis. K.M. performed the bioprinting of the tissues and sectioned all samples. S.S. designed and manufactured the PCB. Y.D. conducted the DIC experiments and analysis. F.S., A.J., and J.H. supported the early development of the mechanical reactor. J.W. and T.T. contributed to the supporting information. L.R. wrote the first draft of the manuscript. M.A.S.S. reviewed the manuscript. M.C. and M.A.S.S. provided supervision and funding.

