## Supplemental file for "Programmable 3D cell alignment of bioprinted tissue via soft robotic dynamic stimulation"

This PDF includes:

Supporting Figure S1: Six-axis low-cost multi-material printer
Supporting Figure S2: Reactor assembly steps
Supporting Figure S3: Printed circuit board design
Supporting Figure S4: Digital image correlation set-up and analysis

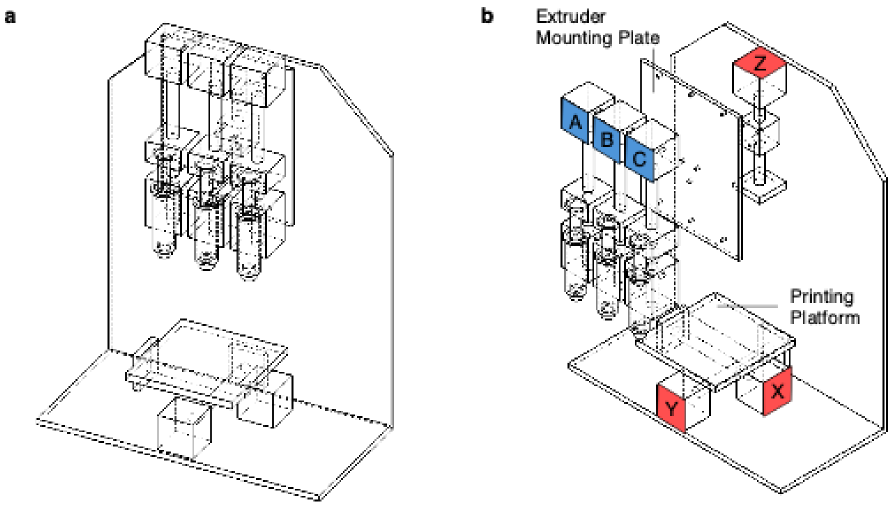


**Fig. S1 | Six-axis low-cost multi-material printer. a,** Drawing of the printer with 3 extruders mounted to a single z-axis driver and xy-plane motorized printing platform. **b,** Details of the ABC extruders fixed to the Z-axis linear motor via the mounting platform and the XY-plane motorized platform with orthogonal linear axis drivers.

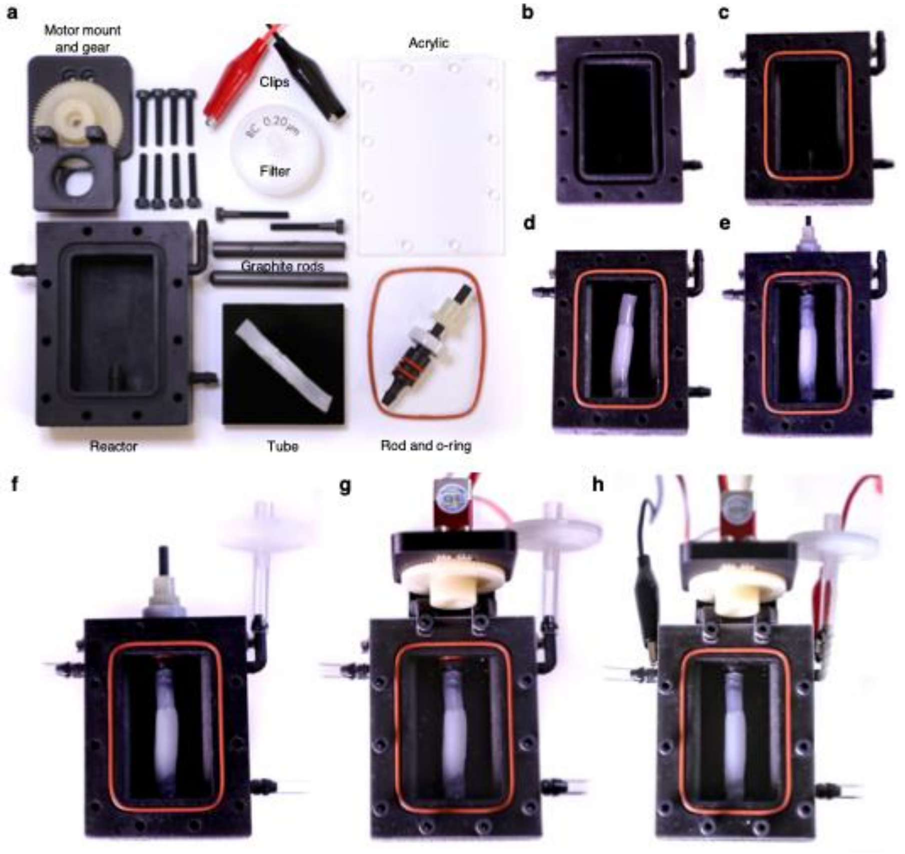


**Fig. S2 | Reactor assembly steps**. **a,** Components of the reactor assembly. **b-c,** Assembly of the mechanical reactor begins with a 3D-printed reactor body **(b)**, where the graphite rods are inserted through the openings in the top, and an o-ring placed into the channel **(c)**. **d-e,** The co-printed silicone-bioink tube is mounted between the barb at the base of the reactor **(d)** and the barb on the rotating rod **(e)**. **f,** The perfusion tubes are connected to the barbs on the left and right, and the sterile filter is connected to the top vent. **g,** After filling the reservoir with media and fixing the acrylic lid, the motor assembly is mounted over the rotating rod and fixed to the reactor body. **h,** Alligator clips are attached to screws at the side of the reactor body, connecting to the graphite rods.

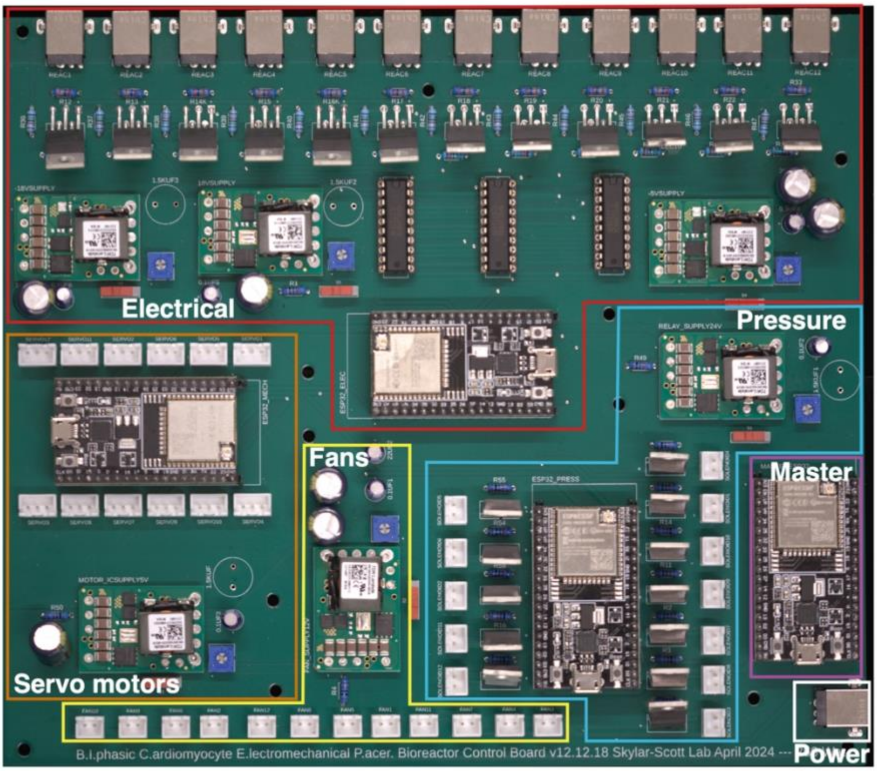


**Fig. S3 | Printed circuit board design**. The image illustrates the components of the board, controlling the electrical stimulation, the servo motors to apply twisting to the constructs, the solenoid manifolds to apply pressure to the constructs, and the fans to prevent the system from overheating. The image further shows the master board and the power connection.

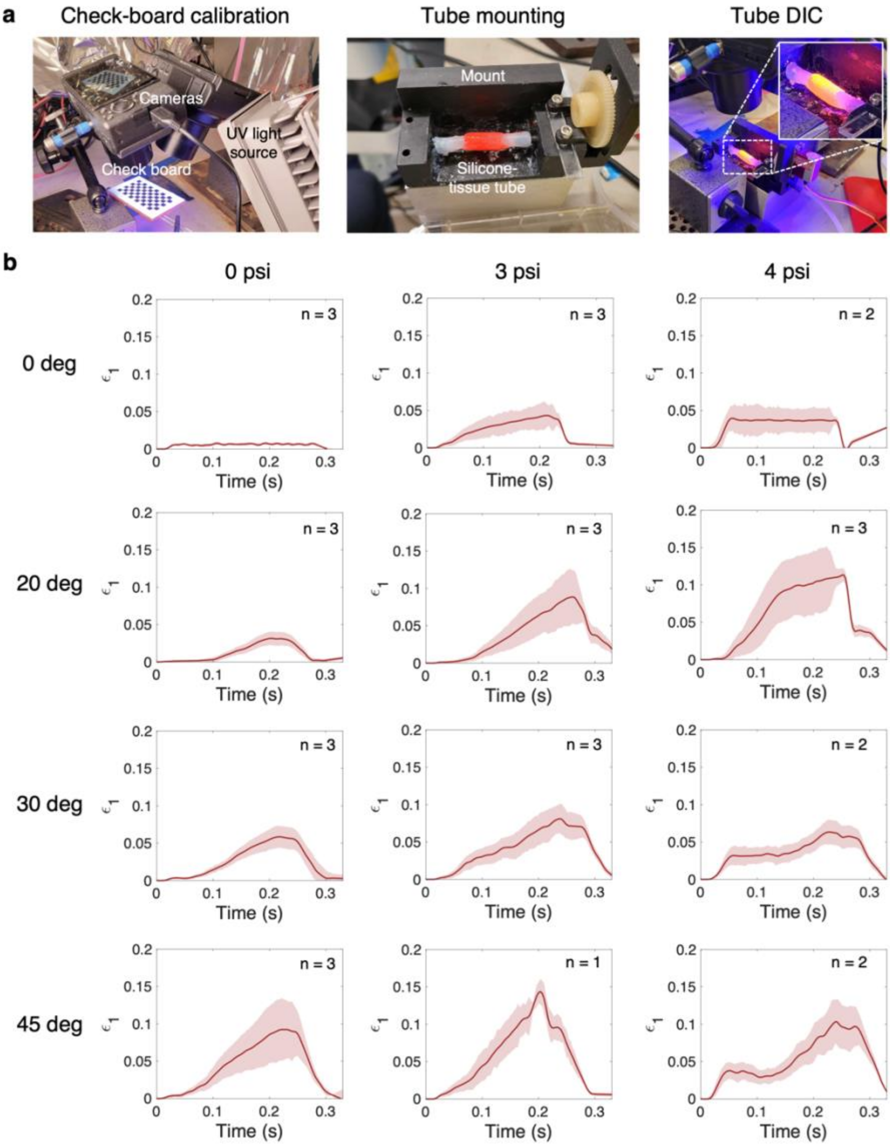


**Fig. S4 | Digital image correlation set-up and analysis**. **a,** Set-up for system calibration, tube mounting, and DIC experiments. **b,** Changes in the first principal strain for 0, 3, and 4 psi expansion, and 0°, 20°, 30°, and 45° twisting angles. All combined conditions are shown.
